# Integrated valorisation of PET and xylose using the oleaginous microorganisms *Yarrowia lipolytica* and *Rhodococcus jostii*

**DOI:** 10.64898/2026.05.30.728943

**Authors:** Alejandro García-Miró, Francisco J. Molpeceres-García, Ines Herrera-Gomez, David Sanz, Alicia Prieto, Jorge Barriuso

**Affiliations:** Department of Biotechnology, Center for Biological Research Margarita Salas, Spanish National Research Council (CIB-CSIC), 28040 Madrid, Spain

**Author notes:** **Corresponding author:** *Jorge Barriuso. Centro de Investigaciones Biológicas Margarita Salas (CSIC), Department of Biotechnology, Ramiro de Maeztu 9, E-28040 Madrid, Spain. Email:.

**Keywords:** *Yarrowia lipolytica*, *Rhodococcus jostii*, terephthalic acid, ethylene glycol, HiC, CalB, mirobial consortia

## Abstract

The increasing accumulation of polyethylene terephthalate (PET) waste has prompted the development of sustainable biotechnological strategies for its degradation and valorisation. This study presents an integrated approach combining enzymatic PET depolymerization by *Yarrowia lipolytica*, engineered to express and secrete the cutinase HiC and the lipase CalB, with the microbial valorization of PET-derived monomers, terephthalic acid (TPA) and ethylene glycol (EG), by *Rhodococcus jostii* RHA1. *Y. lipolytica* was further engineered for xylose metabolism, enabling enzyme production from low-cost lignocellulose-derived substrates. Enzymatic assays with HiC and CalB crudes effectively hydrolysate PET to TPA and EG, demonstrating functional enzymatic activity without purification steps. In addition, *R. jostii* RHA1 was able to use as substrate the released monomers and accumulated intracellular lipids.

Overall, this work demonstrates the feasibility of coupling the production of PET degrading enzymes and microbial lipid, using an abundant monosaccharide, with the assimilation of the PET degradation products to also produce microbial lipids. This modular system provides a promising framework for the sustainable upcycling of plastic waste into value-added bioproducts within a circular economy.

## Introduction

Plastics are versatile and durable materials used to manufacture a huge variety of products. However, their recalcitrance makes them a major environmental problem due to the long period required for their degradation (Yao et al., 2024).

Polyethylene terephthalate (PET) is probably the most popular plastic for single use applications. Its structure is formed by repeated units of terephthalic acid (TPA) and ethylene glycol (EG) linked by ester bonds (Yao et al., 2024). Although only a small fraction of PET waste is currently recycled, there are different available methodologies. Mechanical treatment is the simplest, but the performance of the recycled material is progressively reduced. On the other hand, chemical methods have experienced interesting developments, enabling the recovery of monomers or intermediates that can be repolymerised. However, this approach is not environmentally friendly because of the solvents and the toxic catalysts needed (Qiu et al., 2024; Yao et al., 2024). In contrast, biological approaches offer the possibility of designing clean and efficient processes.

In the last decade, some microorganisms capable of degrading PET have been described. The first one was the bacterium *Ideonella sakaiensis* (Yoshida et al., 2016), that depolymerizes this polymer by the concerted action of two crucial extracellular enzymes: PETase and MHETase. The first one acts on PET releasing the intermediate bis(2-hydroxyethyl) terephthalate (BHET), which is transformed by the same PETase to mono(2-hydroxyethyl) terephthalate (MHET). This compound is degraded into the monomeric units, terephthalic acid (TPA) and ethylene glycol (EG) by the MHETase. It is important to note that the accumulation of MHET can lead to product inhibition of PETase, limiting the overall depolymerization efficiency (Knott et al., 2020).

In addition, the ability of *I. sakaiensis* to metabolize PET-derived monomers is well documented (Yoshida et al., 2016). This capacity has also been described in organisms unable to depolymerize PET. For example, *Comamonas testosteroni* can assimilate TPA (Molpeceres-García et al., 2025) and some strains of *Pseudomonas putida*, as JM37 and GO16, consume EG or both monomers (Molpeceres-García et al., 2026; Orimaco et al., 2025). Similarly, several *Rhodococcus* strains grow on TPA, while *R. jostii* RHA1 can use on both TPA and EG as the sole carbon and energy source (Hara et al., 2007; Shimizu et al., 2024).

Apart of the two enzymes described in *Ideonella*, other microbial hydrolases have demonstrated activity on PET. Among them, cutinases such as TfCut2 secreted by the thermophilic actinomycete *Thermobifida fusca* (Kaur et al., 2023), HiC from *Mycothermus thermophilus* (previously *Humicola insolens*), FsC from *Fusarium solani* (Ronkvist et al., 2009), mostly produce the intermediates BHET and/or MHET. In addition, the lipase B (CalB) from

*Moesziomyces antarcticus* (previously denominated *Pseudozyma antarctica* and *Candida antarctica*) (de Castro & Carniel, 2017) is active on MHET, but not on the polymer and, for this reason, it has been combined with HiC to exploit their synergy for PET degradation (Carniel et al., 2017).

Considering this, several studies have focused on the heterologous expression of PET-degrading enzymes for *in vitro* applications. For example, the PETase and the MHETase from *I. sakaiensis* have been extracellularly produced in two strains of *P. putida*, KT2440 (Brandenberg et al., 2022) and JM37 (Molpeceres-García et al., 2025, 2026). In eukaryotic hosts, the genes encoding the cutinase FsC and some engineered variants have been successfully expressed in *Komagataella phaffi* (Murguiondo et al., 2025), and some modified strains of *Yarrowia lipolytica* can produce CalB (Emond et al., 2010; Park et al., 2019; Theron et al., 2020) or the IsPETase (Kosiorowska et al., 2022). This last yeast is currently one of the most promising chassis in synthetic biology because it is able to produce different metabolites (Miller & Alper, 2019) and to efficiently secrete proteins, has wide metabolic flexibility and advanced genetic toolbox (Madzak, 2018).

In addition, *Y. lipolytica* is capable of producing large amounts of intracellular lipids from diverse and low□cost substrates, including fatty acids and sugars from agro□industrial wastes (Friedlander et al., 2016; Tsirigka et al., 2023) including xylose, the second most abundant monosaccharide in lignocellulosic biomass (De La Torre et al., 2025). This makes it a valuable platform for the bioconversion of lignocellulosic feedstocks into value□added products (Lee et al., 2021). Several studies have focused on enhancing microbial lipid production through molecular and metabolic engineering approaches (Liang & Jiang, 2013).

Within oleaginous organisms, the bacterial genus *Rhodococcus* has also gained increasing attention as non-conventional microbial host due to its remarkable metabolic versatility, robustness, and well-known capacity to accumulate large amounts of intracellular TAGs (Alvarez et al., 2019). In addition, *Rhodococcus* species are also notable for their ability to assimilate aromatic carbon sources derived from environmental pollutants and plastic waste (Solyanikova et al., 2016).

Synthetic biology can contribute to finding innovative solutions not only for the clean degradation of PET, but also for its upcycling. The design of artificial microbial consortia, in which the organisms selected perform complementary tasks to mitigate the accumulation of inhibitory intermediates from PET, such as MHET, may represent a promising approach. Some examples on this have recently been reported, showing the division of labour in the degradation PET□derived intermediates, as reported by Molpeceres-García et al. (2026), may offer a simpler and more robust strategy that minimizes inhibitory effects without extensive system optimization (Qi et al., 2021). Building on this, Crespo-Roche et al. (2025) designed of a bottom□up consortium based on *Y. lipolytica* as microbial chassis to valorise natural polymer, like cellulose and hemicellulose, using two engineered microbial populations.

In this work, we explored an integrated strategy that combines PET depolymerization and valorisation using two oleaginous microorganisms. An engineered strain of *Y. lipolytica* is responsible for PET degradation and lipids accumulation using xylose as substrate, while TPA and EG are upcycled to triacylglycerides (TAG) by *R. jostii* RHA1, highlighting both microorganism as a promising chassis for sustainable upcycling of lignocellulose and PET waste.

## Material and methods

### 1. Media and strains

Yeast and bacterial pre-inocula were grown in YPD and LB media, respectively. The yeast cultures were done in YNB minimal medium without amino acids neither nitrogen (233520, BD Difco™, USA) and NH_4_Cl 0.24 g/L, while M9 minimal medium (Na_2_HPO_4_ 6.78 g/L, KH_2_PO_4_ 3 g/L, NaCl 0.5 g/L, MgSO_4_·7H_2_O 0.25 g/L, FeSO_4_ 9 mg/L, CaCl2 11 mg/L and NH_4_Cl 1 g/L) was used for the bacterium. Vitamins and trace elements solution for the MC medium are described in Barragán et al. (2004). When necessary, amino acids supplements without leucine were added to YNB (Sigma-Aldricht, Germany, REF Y1376). Unless otherwise stated, glucose was the carbon source in the minimal medium. Other carbon sources used were D-xylose (Thermo Scientific, CAS 58-86-6), TPA (Sigma-Aldrich, Ref 185361) or EG (Sigma-Aldrich, Ref 102466).

The *Y. lipolytica* strain used in this work is an obese derivative from Po1d (Le Dall et al., 1994) modified overexpressing the genes *dga1* from *Rhodosporidium toruloides* and *dga2* from *Claviceps purpurea*, inserted in *mef1* locus to increase the lipid yield production (de La Torre et al. 2026). This strain is auxotroph for leucine. The *R. jostii* RHA1 wild type strain was obtained from the German Collection of Microorganisms and Cell Cultures (DSMZ) (Table I).

### 2. Expression of PET-degrading enzymes in Y. lipolytica

Firstly, the gene sequence encoding for the fungal enzymes HiC (ABC06408.1) and CalB (SPO46143.1), were codon-optimized using the Codon Usage Database software (Nakamura et al., 2000) and the sequence from the promoter pTEF (Madzak, 2018) and the signal peptide SP3 (Celińska et al., 2020) were included at the 5′ end of each gene. In addition, a tTEF terminator was used at the 3’ end (Madzak, 2018) of each gene. Both constructs were synthesised by Genescripts (Switzerland) in a pUC vector.

The synthetic construction was cloned in the JMP62 leu integrative vector with the *leu* gene (using the auxotrophy reversion as a marker), using restriction enzymes to construct the plasmid JMP62 HC (Table I). The plasmid contains *Kan*^*R*^, a pBR322 replication origin to replicate in bacteria, and the denominated Z regions (Schmid-Berger et al., 1994) (Supp. Fig. 1).

**Table I.**
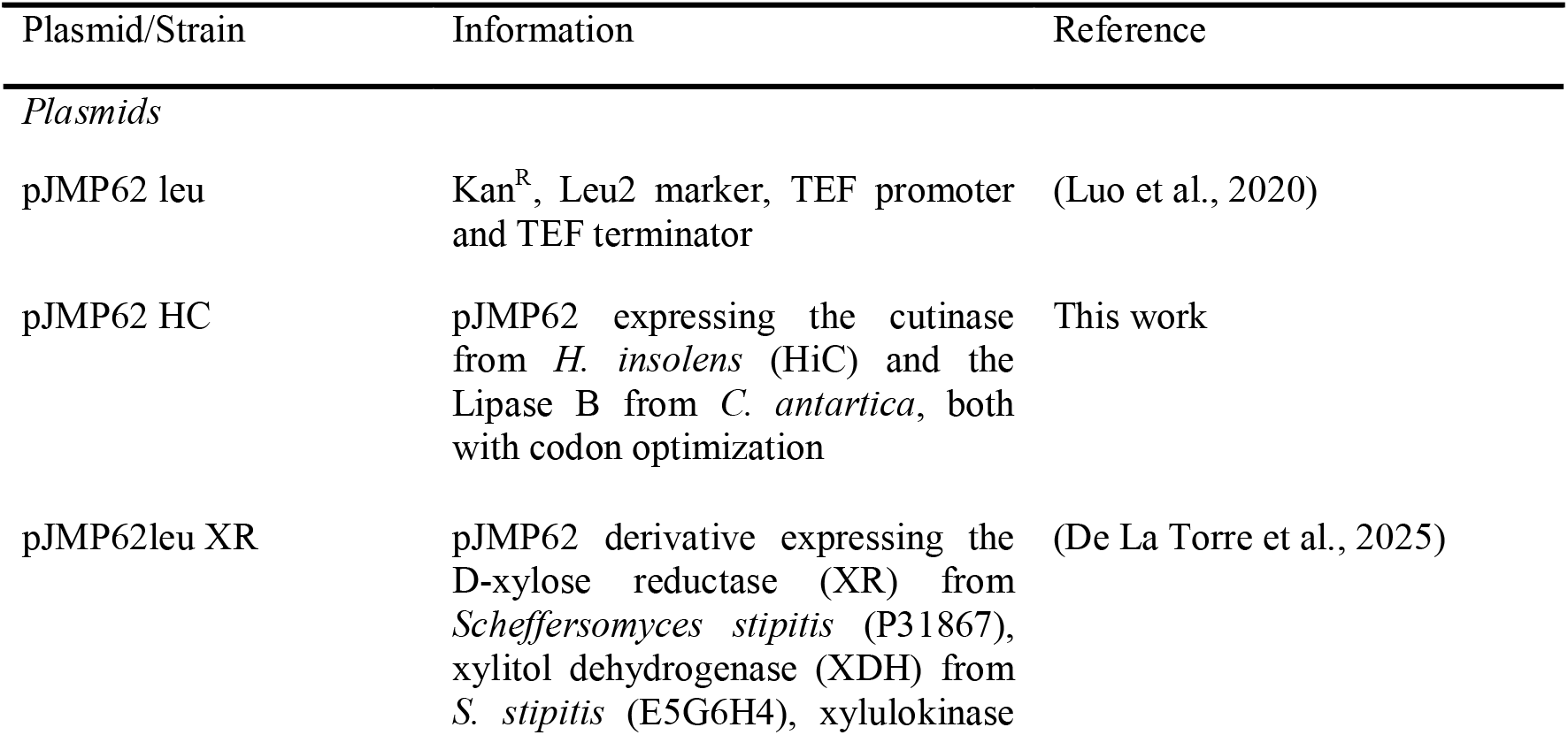
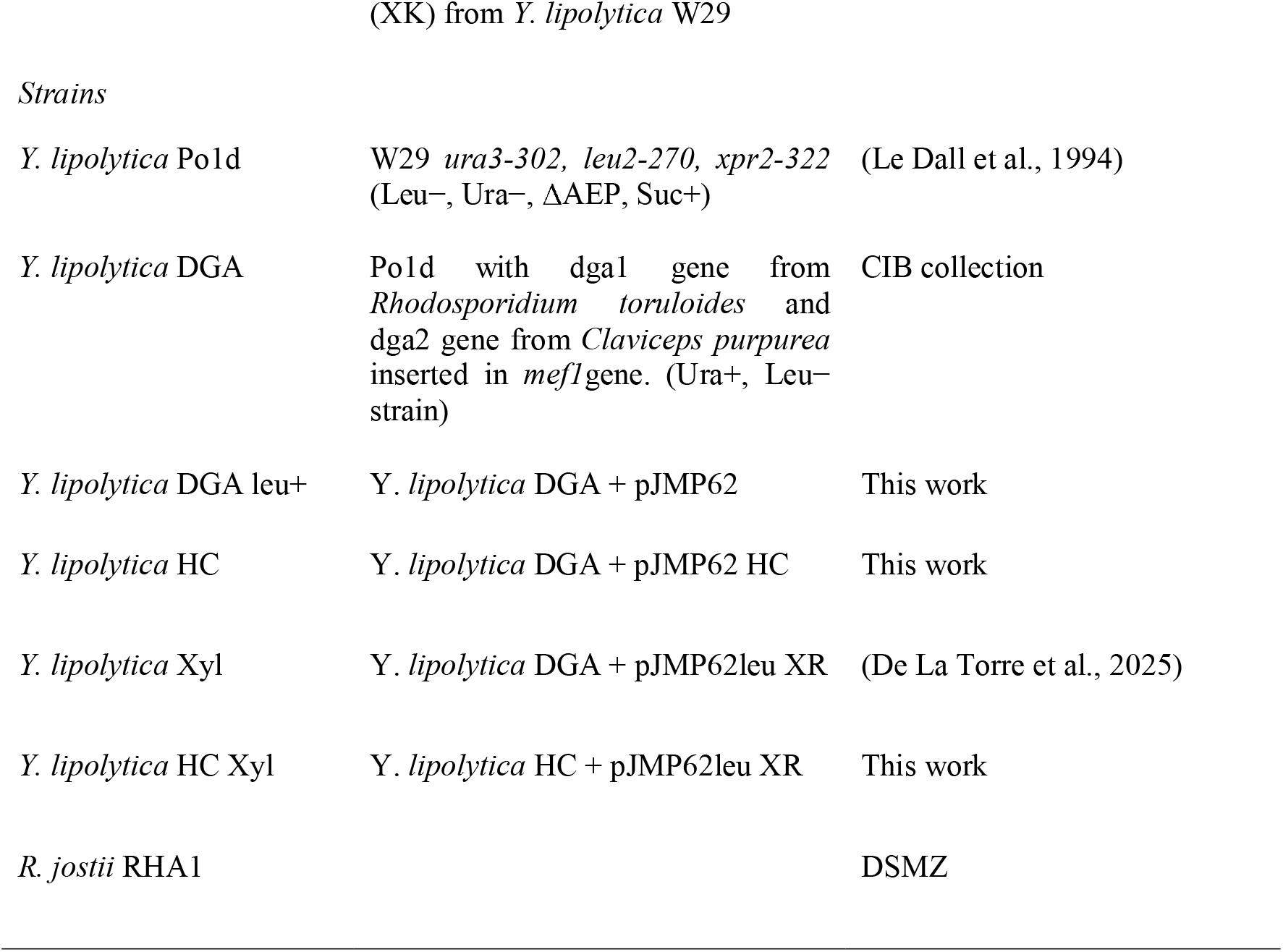
Strains and plasmids used in this work.

In pJMP62 HC both *hiC* and *calB* have their own constitutive promoter, (*pTEF)*, signal peptide SP3 and terminator (*tTEF*). This construct was transformed into the leucine auxotrophic strain of *Y. lipolytica* using the lithium acetate (LiAc) method (Le Dall et al., 1994), and plated in YNB plates with glucose and without leucine. DNA carrier (D9156 Sigma-Aldrich® single-stranded DNA from salmon testes) was used for transformation.

Random clones grown in plates without leucine were selected and PCRs were made to determine the presence of the integrated genes. One primer annealed to the 3′ end of the upstream gene, *hiC*, while the second primer annealed to the 5′ end of the downstream gene, *calB* (Supp. Table 1), allowing amplification only when both genes were present and correctly integrated. After confirming that the genes of interest were inserted in the genome, several clones were screened for unspecific esterase activity using *p*NPB as substrate as in García-Miro et al., (2024). For that, 23 clones were selected from YNB plates without leucine and were grown in a 96-well plate with YPD an o/n. Unspecific activity (*p*NPB) and growth were measured after two days to choose the better candidate. Once assessed the activity of *Y. lipolytica* HC on *p*NPB, the best candidate was transformed with the plasmid pJMP62leu XR to construct the *Y. lipolytica* HC Xyl, and the positives colonies were identified from their ability to grow on xylose plates. The esterase activity against *p*NPB was finally reassessed.

### 3. Enzyme production

Firstly, cultures of 50 mL with the strains *Y. lipolytica* Xyl and HC Xyl growing on xylose were performed, and with *Y. lipolytica* HC Xyl growing on glucose for 3 days with a C/N ratio of 75. Lipids and esterase activity were measured in all the cultures.

To maximize the enzyme production, three 2 L flasks with 400 mL of medium were inoculated at 0.1 OD_600_, *Y. lipolytica* HC Xyl in YNB media without amino acids, with 0.25 g/L of NH_4_Cl and 20 g/L o xylose (ratio C/N 75). After 3 days, the cultures were collected and centrifugated at 11 000 *g* in a Sorvall Lynx 6000 centrifuge equipped with a RF12 rotor (Thermo Fisher). After that, the supernatant was sequentially filtered through 0.8□µm, 0.45□µm and 0.2□µm cutoff membranes (Millipore), and finally the samples were concentrated by ultrafiltration using 10□kDa membranes (Pellicon and Amicon, Merck Millipore, Germany).

The unspecific esterase activity was measured as described above to follow the production of the cultures, and a 12□% PAGE gel with and without SDS was done to determine the presence of HiC and CalB on the concentrated crude due to both of them have activity against *p*NPB, so it was necessary to ensure the production of both enzymes. The commercial HiC (Novozym® 51302, Novozymes) and CalB (Lypozyme ® CalB, Novozymes) were used as controls to determine their migration on gel.

### 4. PET degradation in vitro

2.5 g of amorphous PET (Goodfellow) cut in small pieces were put on bottles of 100 mL with 30 mL of phosphate buffer 100 mM at pH 8. Concentrated enzymatic crude of from section 3 was added at 40 U/mL (against *p*NPB) and then incubated at 50 °C and 250 rpm for 5 days. Every day, unspecific esterase activity and monomer liberation were measured. This experiment was done in triplicate.

The degradation products in the supernatants were analysed by HPLC in an Agilent 1200 series instrument. EG was analysed in a BioRad Aminex HPX-87H column (300 x 7.8 mm) under isocratic flux of 0.5 mL/min of H_2_SO_4_ 2 mM, and detected by refraction index (RID). For BHET, MHET and TPA analysis, a reverse-phase C18 affinity column (ZORBAX Eclipse Plus C18 Agilent Technologies) was used, and the mobile phase consisted of an isocratic gradient of acetonitrile with 0.1% acetic acid and MilliQ H_2_O with 0.1% acetic acid. The gradient changed from 5% of the former to 70% over 20 minutes, with a flow rate of 0.8 mL/min. The temperature was maintained at 30 °C, and the injection volume was 10 μL. The compounds were detected by measuring absorbance at 260 nm using a DAD detector.

Quantification was performed using calibration curves of the standard molecules, analysed under the same conditions at different concentrations.

### 5. *Rhodococcus* growth on TPA and EG

Cultures were done in 24-well plates to determine *R. jostii* RHA1 tolerance to EG and TPA. To do that, a preinoculum was grown in LB medium and the cultures were done in M9 media with 1, 5, 10, 20 and 40 mM of EG, TPA or both. OD_600_ was measured at 6, 24, 32 and 48 h and the cultures were done by triplicate.

Moreover, *Rhodococcus* was fed with the supernatant resulting from the PET degradation, to validate its capacity of growing with PET-released monomers. Therefore, the supernatant containing the PET monomers was filtered by 0.22 µm (REF 12532, Sartorius) under sterile conditions, and the necessary supplements were added to convert the buffered supernatant to M9 media. For that, the MgSO_4,_ NaCl, FeSO_4_ and CaCl_2_ were added, along and the 0.06 g/L of NH_4_Cl to obtain a C/N of 75. The cultures were done for triplicates on 10 mL of volume. The cultures were stop at 24h at which OD^600^ was measured, and the cells were stained with BODIPY to observe the lipid content in the cells on a DM4 B microscope (Leica) equipped with a pE-300 Lite LED illuminator (CoolLED) and a DFC345 FX camera (Leica).

### 6. Lipid quantification

First, for lipid quantification a gravimetric assay was done. For that, the pellets from *R. jostii* RHA1, *Y. lipolytica* HC Xyl, or the consortium were lyophilizated, and 200 mg were use. Then 2.5 mL of HCl 2M was added and, after vortexing 2 min. Sequentially, add 1.8 mL of water and centrifugated 15 mins at 2500 g. The chloroform has to be recovered and placed in a new tube. Add 2 mL of 10% methanol/chloroform and after vertexing again, centrifuge again. This step has to be done twice. Finally, the chloroform is evaporated at 75 °C and then the sample is weighted.

Second, to determine the fatty acids profile of the TAGs produced, fatty acid methyl esters (FAMEs) were done. For that, 20-30 mg of dry biomass was weight, and an internal standard (C21:0, 15 µL at 7 mg/mL) was added. 2 mL of magic methanol were added (methanol/hydrochloric acid/chloroform (10:1:1)). The samples were heated at 90 °C for 1 h, and a washing was done with 1mL of 0.9% NaCl saline solution. Finally, 2 mL of hexane were added, and the samples were centrifugated at 3000 rpm for 3 minutes. The organic fraction was collected and analysed by GC-MS (De La Torre et al., 2025).

## Results and discussion

### 1. Engineering of a recombinant Y. lipolytica strain for xylose utilization and extracellular enzyme production

The first step of this work consisted of constructing a recombinant strain of *Y. lipolytica* that expresses the fungal PET-degrading enzymes HiC and CalB, since they have been reported to act synergistically for full depolymerization of PET (de Castro & Carniel, 2017). HiC exhibits high PETase-like activity and is also capable of catalysing the hydrolysis of MHET, although with limited efficiency, while the high MHETase activity of CalB complements HiC with high efficiency by enhancing the conversion of intermediate products (Carniel et al., 2017).

To obtain the recombinant HC strain, the genes *hiC* and *calB* were expressed in *Y. lipolytica* DGA using the JMP62 vector. Since the integration using this system can occur in different regions of the genome and the number of copies inserted could differ (Gietz & Schiestl, 2007; Le Dall et al., 1994), the esterase activity against the model substrate *p*NPB was measured in several colonies. The values were normalized to OD_600_ reaching a maximum of 87 mU/mL (Supp. Fig 2). Due to the low activity obtained in this assay, it was repeated along 72 h, showing a peak of activity at 72 hours (data not shown). As both enzymes present activity against *p*NPB, the presence of both genes in the selected colonies was further verified via PCR (Supp. Fig. 3).

The second modification consisted on conferring the capability to metabolise xylose to *Y. lipolytica* HC using the plasmid JMP62leu XR (De La Torre et al., 2025). After confirming that the final variant, denominated *Y. lipolytica* HC Xyl, was able to use xylose and produced the expected esterase activity, we tried to identify optimal conditions that balance lipid and enzymatic production comparing its growth on glucose and xylose (Fig. 1). Maximum cell growth in both conditions was observed at 48 h, whereas enzymatic activity continued to increase until 80 h, when the cultures were stopped. Considering the decrease in OD_600_ at this time point, it is unlikely that enzyme production was still active. Moreover, even the enzymatic activity still showed variation at the final time point, it would be incompatible with the lipid accumulation desired because when the carbon source in the medium is depleted, the cell would begin to consume its own reserve lipids. Although slightly lower, no major differences were observed between using xylose or glucose.

**Figure 1.**
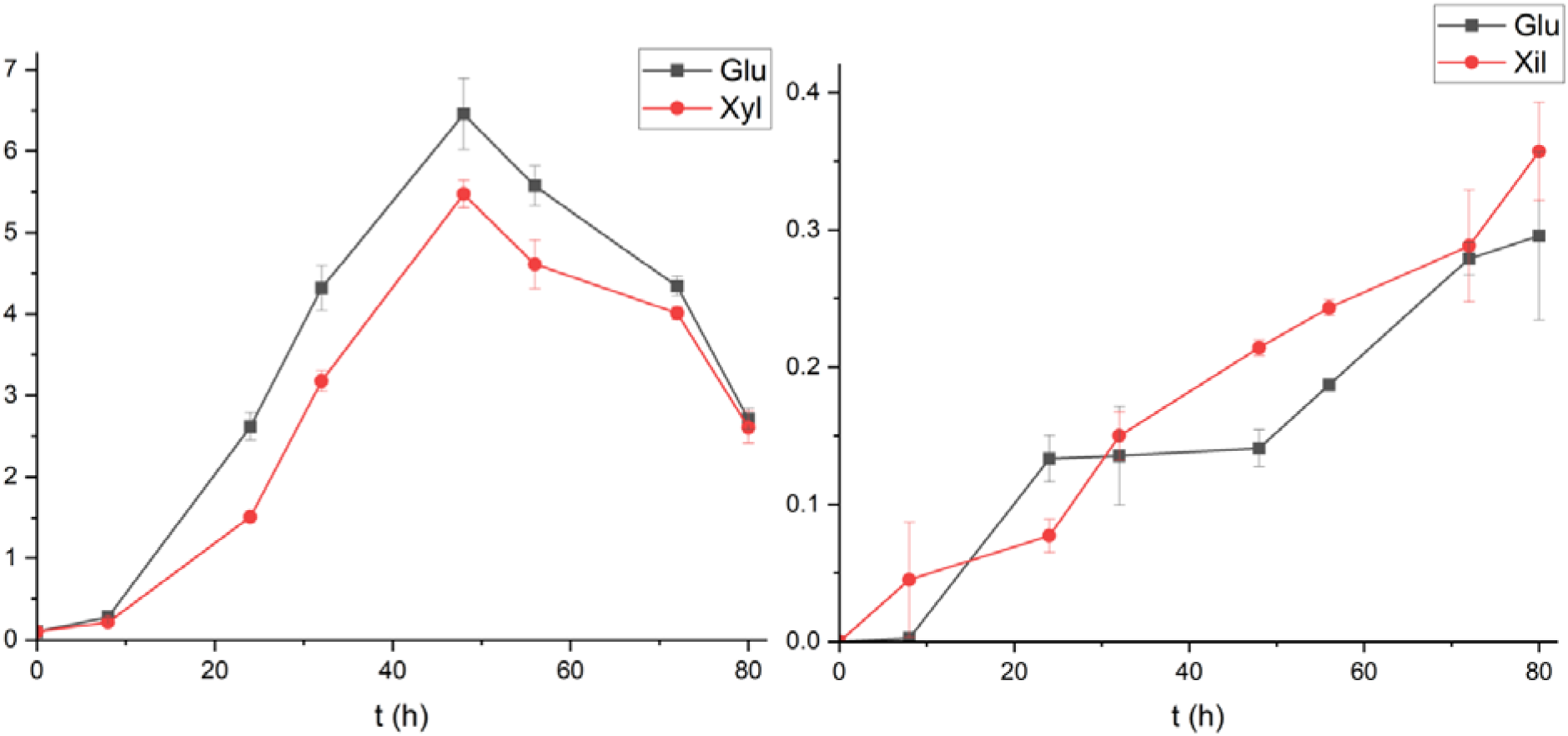
Growth (OD_600nm_) of *Y. lipolytica* HC XylRed in YNB with xylose as sole carbon source (left panel) and activity (U/mL) measured at different times (right panel). 10 g/L of xylose or glucose were used as carbon source and the C/N ratio was set at 75.

### 2. Functional evaluation of Y. lipolytica HC Xyl for enzyme and lipid production

With these preliminary results, further experiments were conducted for three days trying to find a good balance between esterase activity and lipid production in *Y. lipolytica* HC Xyl, grown on glucose or xylose. The strain *Y. lipolytica* Xyl, cultured in xylose, was also tested as a control to evaluate. These conditions were compared to evaluate differences in lipid accumulation and enzymatic production depending on the carbon source and the potential metabolic burden caused by producing the recombinant enzymes.

Gravimetric analyses demonstrated that the strain HC Xyl accumulated 27.4% ± 0.2 lipids, while also producing HiC and CalB, in the medium with xylose. Lipid production is comparable to that obtained with the control strain, *Y. lipolytica* Xyl, which accumulated 27.7% ± 0.4 lipids under the same conditions. This result suggest that enzyme production does not impose an excessive metabolic burden on the yeast. However, when cultivated on glucose, the HC Xyl strain showed significantly higher lipid accumulation (46.1% ± 3.7), likely due to glucose being a more readily metabolizable carbon source.

To grow on xylose, *Yarrowia* requires the expression of 3 heterologous enzymes, a NAD(P)H-dependent D-xylose reductase (P31867) and a xylitol dehydrogenase (E5G6H4) from *Scheffersomyces stipites*, and the xylulokinase (YALI0F10923) from *Y. lipolyitca* W29 (De La Torre et al., 2025), previously cloned into the strain. Although the associated metabolic stress seemed to be relatively low, this strain was expected to grow worse in xylose than in glucose due to intrinsic metabolic and regulatory limitations, including low flux through of the native xylose pathways, cofactor imbalances, and bottlenecks such as inefficient xylitol dehydrogenase activity (Zhong et al., 2025). Nevertheless, xylose metabolism proceeds through the pentose phosphate pathway (PPP), which is a major source of NADPH in *Y. lipolytica*, a key reducing equivalent required for fatty acid synthesis (Lazar et al., 2018).

Regarding nitrogen limitation, Supp. Fig. 5 shows that a C/N ratio of 75 boost lipid accumulation meanwhile the enzymes of interest were produced. This represents a medium-high C/N ratio (Kuttiraja et al., 2016).

For further enzyme production the yeast was cultured in 2□L flasks under the same conditions, recovering 0.8 L of the supernatant. After confirming enzyme expression by PAGE (Supp. Fig. 4) and extracellular esterase activity against *p*NPB (0.74 U/mL), the culture supernatant was concentrated to a final volume of 2.5 mL with 232.3 U/mL of activity and stored for *in vitro* experiments. The growth and enzymatic production under these conditions were slightly better to those observed in smaller cultures (Fig 1). The increased volume and better oxygenation could have resulted in better growth and enzymatic production.

It is important to note that, even though the genes that encode HiC and CalB are placed in the same genomic region and have the same promoter, signal peptide and terminator, the intensity of the bands observed in the gel is slightly different (Supp. Fig. 4). Although this phenomenon has been extensively studied in yeasts (Aza et al., 2021), particularly in *Y. lipolytica*, it is well established that signal peptide efficiency is strongly dependent on the target protein. Different signal sequences can lead to variable secretion performance depending on the protein context, highlighting the need for case-by-case optimization (Celińska et al., 2018). For a better trade-off between lipid and enzymatic production, optimization could be carried out in larger bioreactors.

### 3. Assessment of in vitro degradation of PET film with the enzymatic cocktail produced by Y. lipolytica HC Xyl

*In vitro* degradation of PET film was done by adding 40 U_*p*NPB_/mL of the enzyme cocktail produced by *Y. lipolytica* to the reaction mixture. Enzyme activity was monitored every day during incubation, detecting its gradual decrease (data not shown) and, at day 5, when the reactions were stopped, there was no residual activity. At the final time, the TPA released amounted to 8.7 ± 2.4 mM and no MHET or BHET were detected (Fig. 2). This suggests that, as expected, the first degradation step, PET conversion into BHET, is the limiting step of the reaction. Carniel et al. (2017) reported a total release of 2.23 mM monomers from pre-treated PET in 14 days using 20 mg g□^1^ of a HiC-CalB enzyme mixture (9:1), which represented their best result. The hydrolysis products consisted mainly of TPA (62%), followed by MHET (35%), with negligible amounts of BHET. In their study, reaction parameters such as temperature, pH, buffer system, and enzyme ratio were optimized for PET degradation. In the present work, similar reaction conditions were applied, with the exception of enzyme concentration and PET substrate. Regarding enzyme stability, previous studies have reported measurable PET-degrading activity persisting for two weeks or longer when using commercial enzymes (Carniel et al., 2017; Eugenio et al., 2022), which did not occur in our case. Enzymatic stability is strongly influenced by numerous parameters (Eugenio et al., 2022), and commercial enzyme formulations typically contain stabilizing agents (Ortiz et al., 2019) designed to preserve enzymatic activity over extended periods like the immobilized CalB used in Carniel et al., (2017), which may account for the differences observed in reaction time and activity profiles compared with our system.

**Figure 2.**
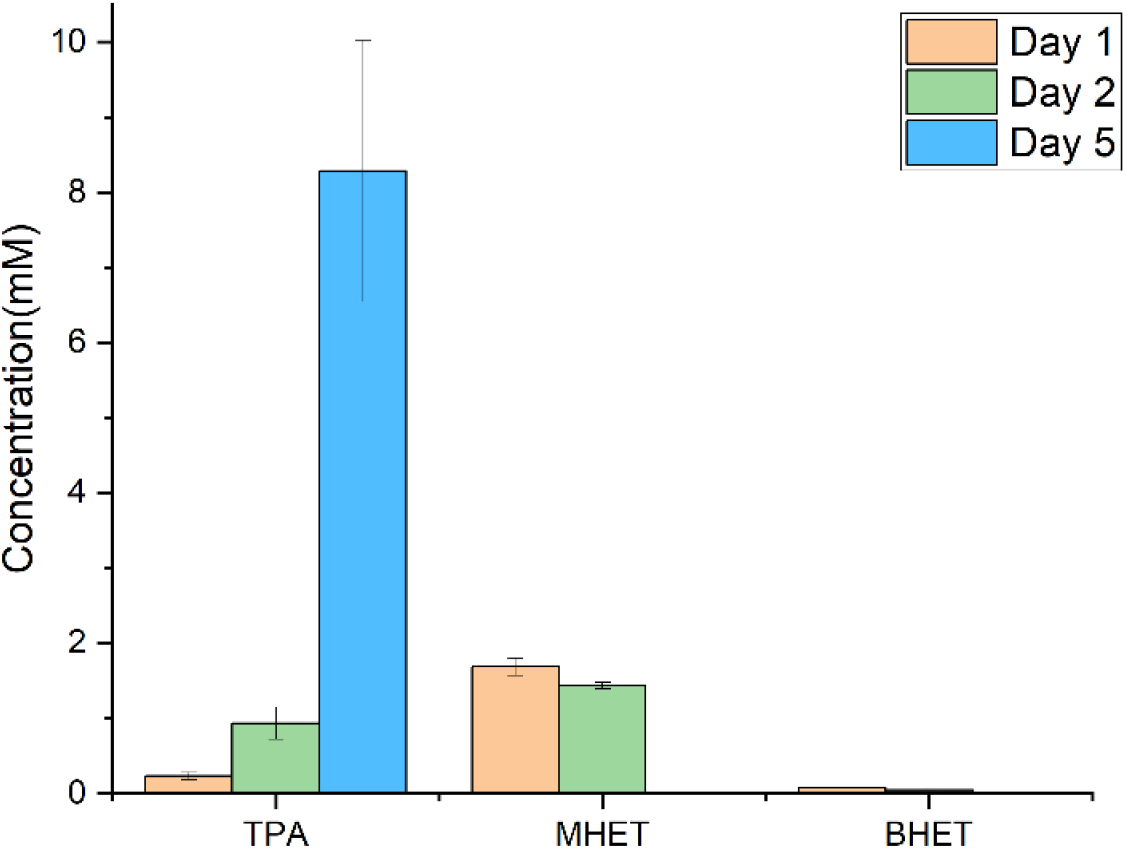
Products released after *in vitro* hydrolysis for 5 days with 2.5 g of PET film and the concentrated supernatant of *Y. lipolytica* HC containing 40 U_*p*NPB_ /mL.

Eugenio et al., (2022), treated different amounts of post-consumer PET with 1□mg·mL□^1^ of HiC, resulting in substantial monomer release that correlated with PET loading. In the condition most comparable to ours, approximately 80□g·L□^1^ of PET, a final monomer concentration of around 50□mM, was detected at the reaction endpoint. Other studies describe the use of the FAST-PETase, an improved version of the IsPETase (Lu et al., 2022), reporting total depolymerization of PET films with 200 nM of pure enzyme.

Taken together, the previously discussed results indicate that PET degradation generally requires the use of purified or immobilized enzymes at relatively high loadings. However, enzyme purification typically entails considerable time investment and substantial costs, which may limit the practical applicability and scalability of such systems. Although the approach presented in the current work requires further optimization to enhance PET depolymerization efficiency, it offers a viable scheme that links plastic waste degradation with the sustainable production of value-added lipids and bioproducts using a robust microbial platform within an integrated circular economy framework.

### 4. Identification of Rhodococcus jostii RHA1 as a chassis for lipids production from EG and TPA

As previously mentioned, bacteria of the genus *Rhodococcus* can be used for the biorecycling of PET monomers. Several studies have documented the mechanisms of *Rhodococcus* species to assimilate EG (Shimizu et al., 2024) and TPA (Hara et al., 2007). Here, we worked with *R. jostii* RHA1 due to its ability to utilize both EG and TPA, unlike other *Rhodococcus* species that can only metabolize TPA, as well as its strong oleaginous capacity.

Different concentrations of TPA and EG were tested, independently or simultaneously, to assess their potential effect on *R. jostii* RHA1 growth (Fig. 3). In general, the combination of TPA and EG (Fig. 3C) seemed to be positive for the bacterium within the concentration range tested. This is expected as the combined substrate conditions also contained a higher total amount of available carbon compared to the other condition. The highest OD_600_ value was observed in the presence of both monomers at 40 mM each, while at the highest concentration (50 mM) there was a slight inhibition. All cultures behave similarly with 1 mM substrate, showing very low bacterial growth. A maximum was reached at 24 h and then growth remained stable or slightly decreased. Similarly, at the highest concentration tested (50 mM), all cultures showed a subtle decrease in OD_600_ after 48 h when compared to the values observed with 40 mM.

**Figure 3.**
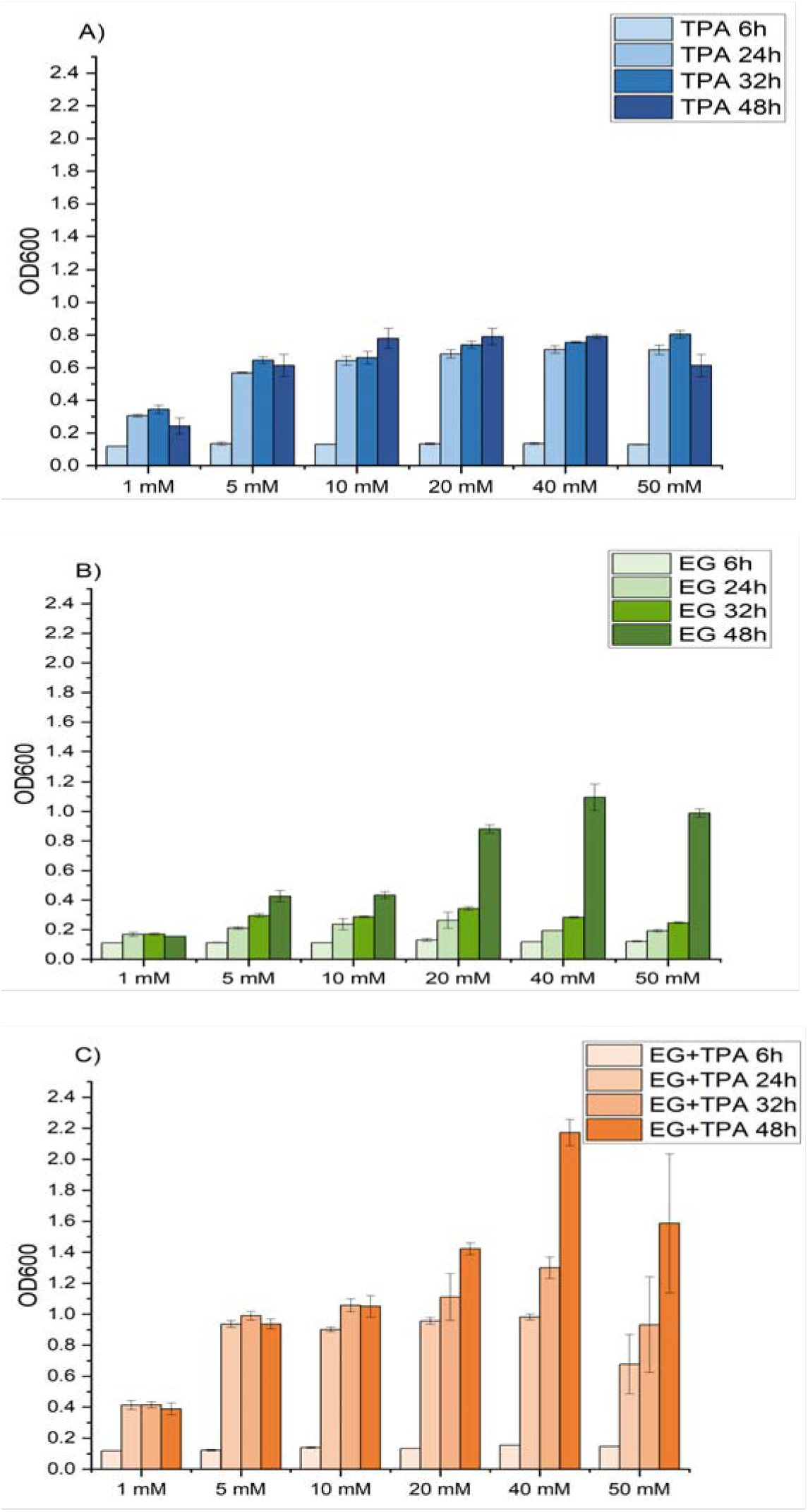
Growth of *R. jostii* RHA1 cultures of 48 h in media containing 1, 5, 10, 20, 40 and 50 mM of: A) TPA, B) EG or C) both together.

When using only EG (Fig. 3B), the OD_600_ values remained stable over the first 36 h, followed by an increase at 48 h at concentrations between 20-50 mM. With only TPA as the carbon source (Fig.3A), the maximum growth was also achieved at 40 mM, despite no significant differences were observed with 50 mM, this indicates that the TPA at high concentrations can produce certain toxicity. It should be noticed that these assays were performed in microplates, where oxygen availability is reduced, while the dioxygenases responsible for the metabolic process of TPA uptake have high oxygen demand (Hara et al., 2007). To overcome this problem, we tested bacterial growth in flasks with 20 mM of TPA, achieving a much larger OD_600_ value (3.10 ± 0.36).

Finally, once demonstrated the capacity of *R. jostii* RHA1 to utilize TPA and EG, the products obtained through enzymatic hydrolysis of PET films with HiC and CalB were used as a carbon source. The supernatant resulting from this reaction was conditioned into M9 medium and inoculated with *R. jostii* RHA1. After 24 h, an OD_600_ of 0.88 ± 0.01 was observed, confirming the bioavailability and assimilation of the released monomers. These results suggest that such compounds can be metabolized to support growth and lipid production (Supp. Fig. 6), pointing to the potential of this approach for PET upcycling applications.

### 6. Lipid profile

It is well established that the nature of the substrate can influence the fatty acid profile accumulated by oleaginous microorganisms (Castro et al., 2016; Fabiszewska et al., 2019). In *Y. lipolytica*, lipids can be synthesized via two pathways: a *de novo* route using non-lipid carbon sources such as sugars, and an *ex novo* route involving direct fatty acid uptake from the medium (Fabiszewska et al., 2019). According to Ledesma-Amaro et al. (2016) approximately half of the fatty acids released from the triglycerides accumulated by *Y. lipolytica* grown on xylose corresponded to oleic acid (C18:1), followed by palmitic (C16:0), stearic (C18:0), and linoleic acid (C18:2). In contrast, lipid metabolism in *R. jostii* RHA1 has been less extensively characterized. However, it is known that this strain can synthesize lipids from aromatic substrates (Dean-Ross et al., 2001). Xiong et al. (2016) reported a lipid profile dominated by palmitic (C16:0), and with abundance of heptadecanoic (C17:0) and heptadecenoic (C17:1) acid and minor amounts of palmitoleic (C16:1), oleic (C18:1), linolenic (C18:3), and stearic acid (C18:0), and even a little of pentadecanoic (C15:0). This profile is very similar to the one identified in the current work (Supp. Fig. 7) but in contrast to Xiong et al. (2016), we observed linoleic acid (C18:2).

## Conclusions

This study demonstrates the potential of combining enzymatic PET depolymerisation with microbial lipid production in an integrated biotechnological platform. The engineered *Y. lipolytica* HC Xyl strain was able to grow on xylose, simultaneously producing PET-degrading enzymes (HiC and CalB) and accumulating intracellular lipids without a significant compromise in lipid yield. These characteristics highlight its suitability as a dual-purpose chassis for both enzyme secretion and bioproduction from low-cost substrates.

The crude enzymatic extracts produced by *Y. lipolytica* were effective in depolymerising PET, yielding TPA and EG as the main products. Although the efficiency of this crude cocktail was lower compared to systems based on purified or immobilized enzymes, this approach offers a clear advantage in terms of process simplicity, cost reduction, and scalability. Importantly, the generated monomers were readily assimilated by *R. jostii* RHA1, supporting its role as a robust organism for the valorisation of PET-derived compounds into lipids.

Together, these results support the feasibility of a two-stage or modular system in which PET is first enzymatically depolymerised and subsequently converted into value-added products by specialized microorganisms. Future work should focus on optimizing enzyme production, improving PET hydrolysis efficiency, and refining microbial consortia strategies to enable one-pot processes and enhance overall process performance. This integrated platform exemplifies a sustainable strategy linking enzymatic PET degradation with microbial lipid biosynthesis, advancing the circular bioeconomy by transforming plastic and xylose waste into valuable bioproducts.

## Acknowledgements

This work was supported by the Spanish project MOLA (PID2024-162673NB-I00 MICIU/AEI/10.13039/501100011033 and European Union). The authors acknowledge the support toward the publication fee by the CSIC Open Access Publication Support Initiative through its Unit of Information Resources for Research (URICI).

## Notes

### Competing Interest Statement

The authors have declared no competing interest.

