## Supplementary Material for "Integrated valorisation of PET and xylose using the oleaginous microorganisms *Yarrowia lipolytica* and *Rhodococcus jostii*"

Supp. Table 1. Pimers used to determine the insertion of *hiC*and *calB* on *Y. lipolytica* DGA.

| Primer name | Sequence | Tm |
| --- | --- | --- |
| HiC Fw 3 | 3’-TTTGCCCGaGGAAGCACCG-5’ | 61 |
| CalB Rv 3 | 3’-GGGTCACGATTCCGCTACA-5’ | 58 |

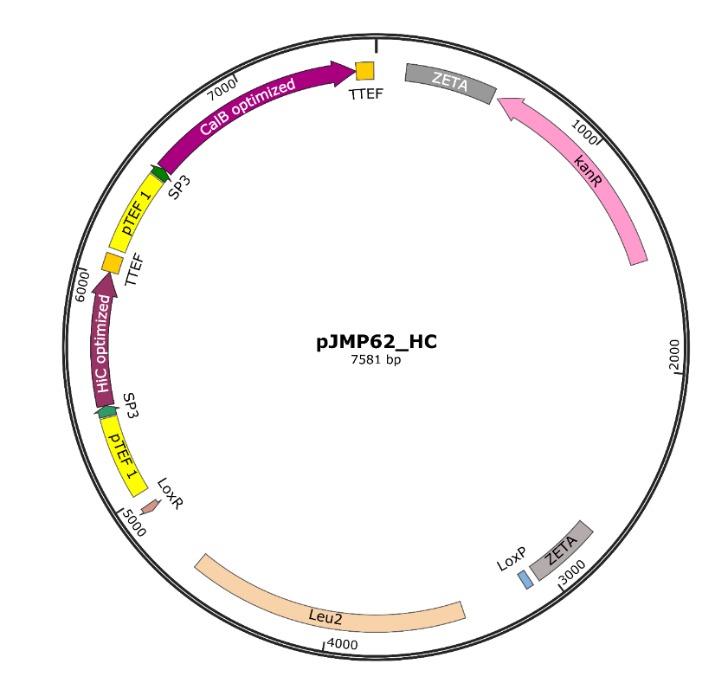

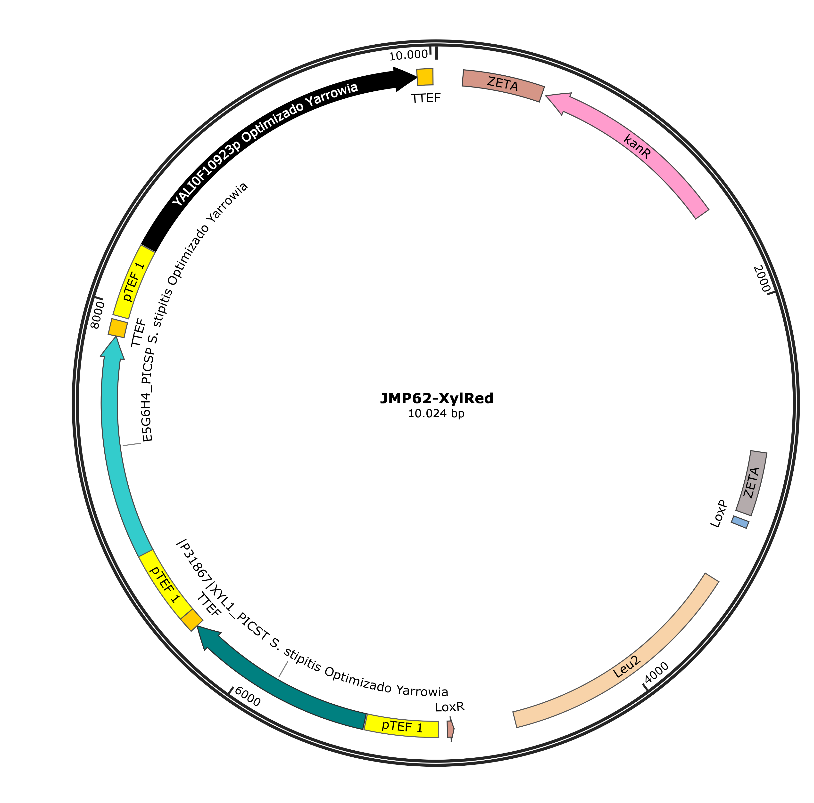

**Supp. Figure 1**. Plasmid maps used in this work for (A) HiC and CalB production, or for (b) xylose assimilation.

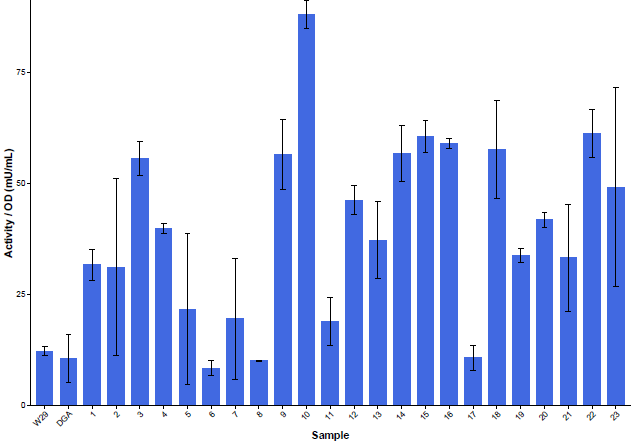

**Supp. Figure 2.** Activity against *p*NPB.of *Y. lipolytica* HC transformants after an o7n growing in YPD in a 96-well plate. The different colonies were grown and tested by triplicates, and the wild type (W29) and the strain modified to accumulate lipids were used as negative controls.

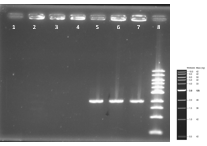

**Supp. Figure 3**. Agarose gel at 0.7%. The 1-3 samples are PCRs from *Y. lipolytica* DGA and the 5-7 from *Y. lipolytica* HC with the primers HiC Fw (3’-TTTGCCCGaGGAAGCACCG-5’) and CalB Rv (3’-GGGTCACGATTCCGCTACA-5’). Each PCR was done at three different TM, 57, 59 and 61 °C. The amplifications only occurred in the strain *Y. lipolytica* HC in which the genes *hiC* and *calB* were integrated.

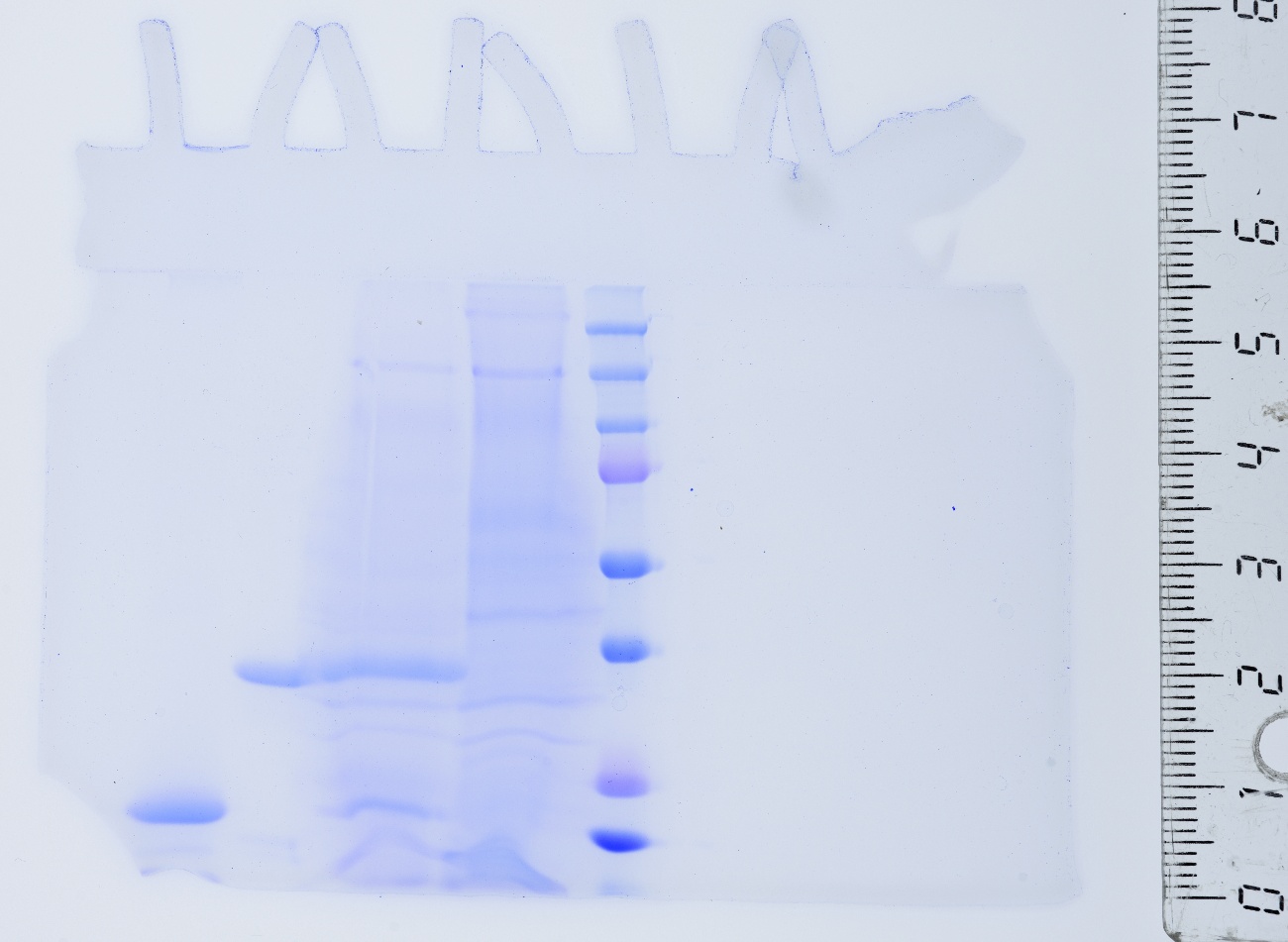

**Supp. Figure 4.** Acrylamide gel at 12%. The samples are, from left to right, HiC (Novozym® 51302, Novozymes), and CalB (Lypozyme ® CalB, Novozymes), *Y. lipolytyca* DGA, *Y. lipolytica* HC and the protein standard.

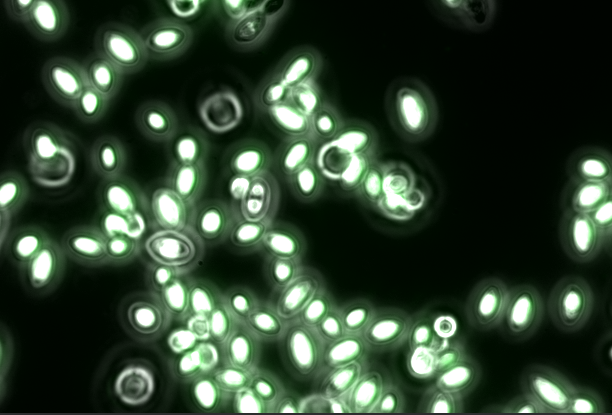

**Supp. Figure 5.** *Y. lipolytica* HC or *Y. lipolytica* HC XylRed after 72h grown in glycerol (A) or xylose (B) stained with bodipy X g/mL.

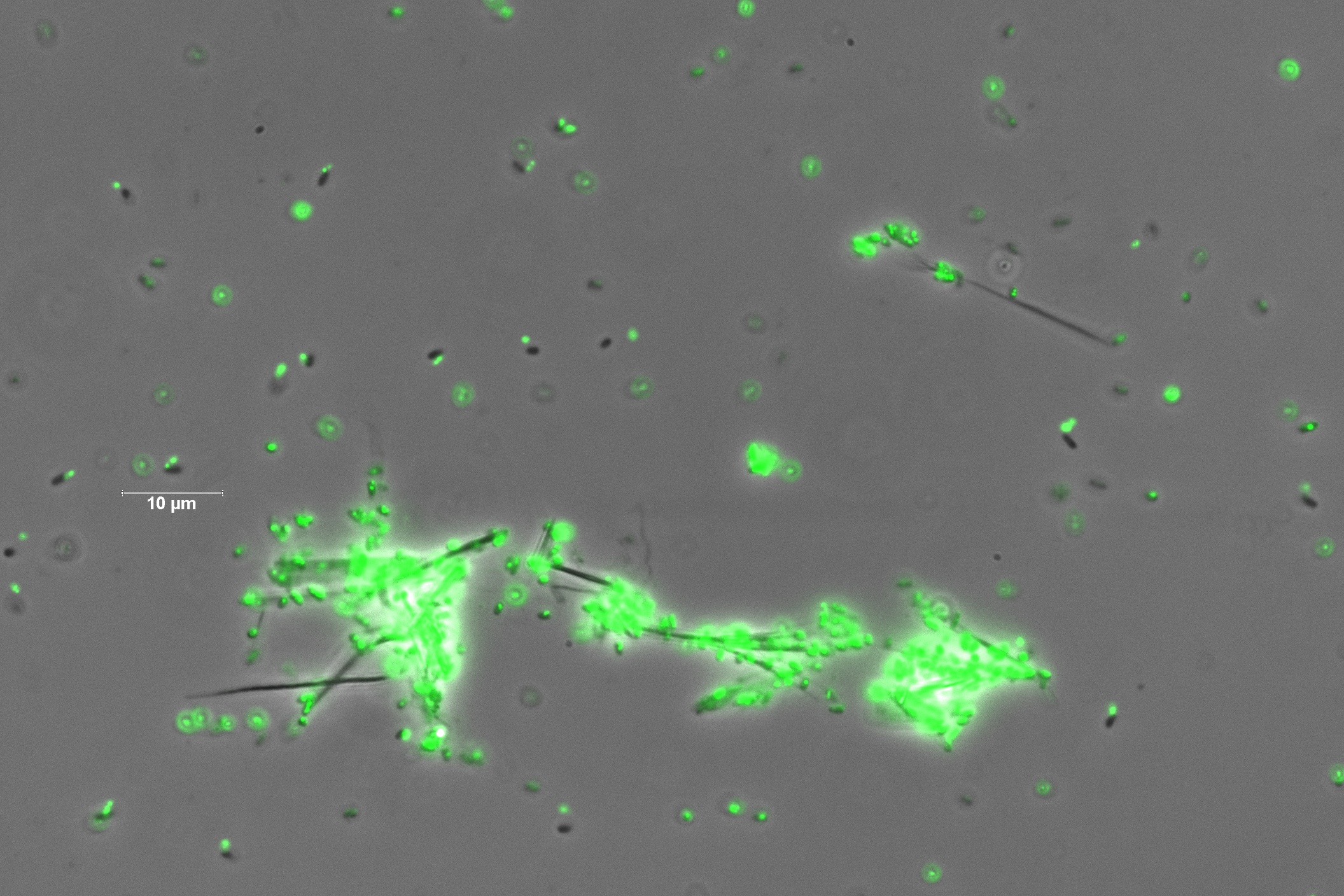

**Supp. Figure 6.** *R. jostii* RHA1 after 24h growing in the supernatant resulting from the PET enzymatic degradation and transformed into MC media. The cells have been stained with bodipy X mg/mL.

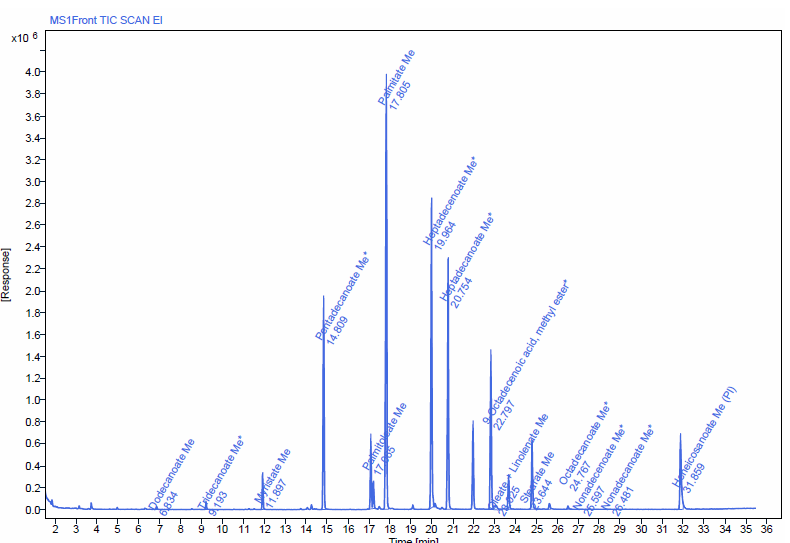

**Supp. Figure 7**. LC–MS chromatogram of lipids derivatized to FAMEs from *R. Jostii* RHA1 growing on TPA. Peaks correspond to individual FAMEs identified based on retention time and mass spectral characteristics
